# Impedance-Derived Heart Rate and Heart Rate Variability from tDCS Output Voltage: Sensorless Physiological Monitoring During Electrical Neuromodulation

**DOI:** 10.64898/2026.07.29.741585

**Authors:** Yahia Abdalla, Benjamin Babaev, Kevin Walsh, Catarina Ferraz, Leigh Charvet, Giuseppina Pilloni, Marom Bikson, Mohamad FallahRad

## Abstract

**Background:** Transcranial direct current stimulation (tDCS) devices adjust output voltage to maintain the target current despite varying impedance. Pulsatile blood flow produces beat-synchronous changes in tissue impedance.

**Objective:** To determine whether impedance-derived heart rate (IHR), heart rate variability (IHRV), and respiration (IDR) can be estimated from tDCS output voltage without additional physiological sensors.

**Methods:** A custom analog front-end acquires the tDCS output voltage across its full dynamic DC range and superimposed AC fluctuations with high precision. Beats detected from the AC-coupled signal yielded normal-to-normal intervals for HR, HRV, and interval-derived respiration. Accuracy was quantified as mean absolute error (MAE) in 10 healthy laboratory participants against ECG and respiration-monitor references, and against chest-strap RR intervals in 19 at-home sessions from 10 participants with mild-to-moderate depression.

**Results:** Laboratory MAEs versus ECG were 0.57 bpm for HR, 9.40 ms for SDNN, and 18.90 ms for RMSSD (r = 0.995, 0.859, and 0.778; N = 10); respiratory-rate MAE was 1.36 breaths/min (r = 0.852; N = 6). Across 1-5 mA of tDCS, the cardiac voltage ΔV_cardiac_(t) amplitude scaled linearly with current (slope, 0.073 mV/mA; p < 0.001). The pulsatile impedance ΔZ_cardiac_(t) = (ΔV_cardiac_(t)/I_applied_) amplitude averaged 0.080 ± 0.029 Ω (mean ± SD) across 50 participant-current observations, with no significant dependence on intensity (slope, −0.002 Ω/mA; p = 0.086). At-home MAEs were 1.43 bpm for HR, 8.86 ms for SDNN, and 24.92 ms for RMSSD (r = 0.995, 0.767, and 0.680; 19 sessions).

**Conclusions:** tDCS output voltage contains a recoverable cardiac-synchronous signal arising from pulsatile impedance, enabling HR, HRV, and respiratory monitoring without additional physiological sensors.

## 1. Introduction

Transcranial direct current stimulation (tDCS) delivers low-intensity (few mA) direct current through scalp electrodes [1]. Current-controlled devices adjust output voltage to maintain the target current as stimulation-path impedance varies between subjects and over time [1-3]. Pulsatile blood flow can produce small beat-synchronous impedance changes that are measurable with specialized impedance plethysmography and impedance cardiography equipment [4,5]. We hypothesized that heartbeat-related impedance changes in the stimulation path would appear as small cardiac-synchronous voltage fluctuations in the tDCS stimulator output.

The stimulator output voltage can be described as the sum of the impedance-dependent voltages that scale with applied current and additive voltage terms not generated by the stimulation current:

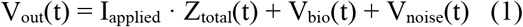

where I_applied_ is the applied current (A), Z_total_(t) is the stimulation-path impedance (Ω), V_bio_(t) represents endogenous biopotentials such as ocular and myogenic activity (V), and V_noise_(t) is instrumentation/environmental noise (V). We decompose the total impedance Z_total_(t) into a baseline term and three time-varying components (in Ω):

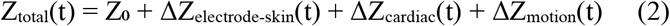

Z_0_ is the time-invariant impedance. ΔZ_electrode-skin_(t) represents changes at the electrode/skin - a significant source of impedance change during transcutaneous stimulation [6], typically observed as a gradual impedance drift during tDCS [3]. ΔZ_cardiac_(t) represents heartbeat-related impedance changes [4,5]. ΔZ_motion_(t) is the impedance change associated with electrode motion artifacts [7]. Substituting Eq. (2) into Eq. (1) yields:

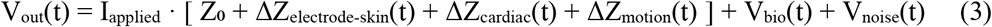

Accordingly, the cardiac-synchronous impedance contribution to the stimulator output voltage is:

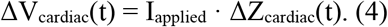

We aimed to extract ΔV_cardiac_(t) from the stimulator output to identify normal-to-normal (NN) interbeat intervals and derive cardiac and respiratory metrics. Specifically, we hypothesized that the ΔV_cardiac_(t) component of the tDCS output voltage is recoverable for estimation of impedance-derived heart rate (IHR), impedance-derived heart rate variability (IHRV), and impedance-derived respiration (IDR). IDR is derived from the impedance-based beat-to-beat intervals, analogous to ECG-derived respiration [8].

While the biophysics of our IHR approach is conceptually analogous to impedance plethysmography (IPG) and impedance cardiography (ICG) [4,5], IPG/ICG apply high-frequency, low-amplitude currents through dedicated source and sensing electrodes, whereas the present approach uses the tDCS current itself and measures voltage across the same stimulation electrodes. Studies combining transcranial electrical stimulation (tES) with EEG or MEG report heartbeat-related nuisance artifacts [9-11]. Here, we engineer a system in which cardiac-synchronous voltage fluctuations in the tDCS device’s output can accurately detect physiological signals.

Heart rate (HR), heart rate variability (HRV), and respiratory rate are general biomarkers of physiological state and autonomic regulation [12,13], and tDCS itself has been reported to affect HRV [14]. During stimulation, these measures are relevant both as physiological outcomes and as candidate inputs for adaptive neuromodulation. Acquiring them currently requires separate sensing hardware such as ECG electrodes, a respiration monitor, or a photoplethysmography sensor. Monitoring them without additional sensors or electrodes would therefore broaden the use of physiological measurement across tDCS applications, including home use [15].

We present the first implementation and validation of this approach. A custom analog front-end recorded the tDCS output voltage during laboratory validation at 2 mA (Experiment 1), a 1–5 mA current sweep (Experiment 2), and remotely supervised home tDCS (Experiment 3). We characterized the cardiac-synchronous component across experiments, evaluated its scaling with applied current, and validated the resulting HR, HRV, and respiratory-rate estimates against cardiac and respiratory references.

## 2 Methods

### 2.1 Participants

#### In-lab cohort (Experiments 1 and 2)

Ten healthy adults (5 female, 5 male; age range 21–54 years; mean ± SD, 30.7 ± 12.9 years) participated. The study was approved by the Institutional Review Board (IRB) of The City College of New York (protocol 2024-0013-CCNY), and all participants gave written informed consent.

#### At-home cohort (Experiment 3)

Ten unmedicated adults with mild-to-moderate depression (4 female, 6 male; age range 36–56 years; mean ± SD, 45.6 ± 5.6 years) participated in an ongoing remotely supervised at-home tDCS study. The study was approved by the IRB of NYU Langone Health (protocol s24-00536; registered as NCT06455527), and all participants gave written informed consent. At-home sessions were selected for complete logger recordings and a synchronized Polar H10 reference.

### 2.2 Stimulation hardware and recording setup

In the in-lab cohort, stimulation was delivered by a low-noise constant-current source [16] paired with high-capacity tDCS (HC-tDCS) hydrogel electrodes [17]. Electrodes were placed in a bifrontotemporal stimulation montage, with the anode positioned over the left eyebrow and the cathode over the right eyebrow.

A custom analog front-end recorded the tDCS output voltage without perturbing the delivered current (Supplementary Figure S1). The AC-coupled cardiac path comprised a 0.5 Hz first-order high-pass stage, protection circuitry, a 100× amplifier, a 33.8 Hz 5th-order elliptic low-pass anti-aliasing stage, and a 17.44× output gain stage. Together, these stages gave a nominal gain of 1744× and produced the AC-coupled signal used for IHR detection. A parallel DC-coupled path used a high-impedance attenuator (0.14×), protection circuitry, and a low-noise, high-impedance buffer to preserve the slowly varying baseline output voltage.

In-lab signals were acquired at 1 kHz with an ADInstruments PowerLab 8/35 (ADInstruments, Sydney, Australia; Figure 1B). Up to five channels were recorded per session: single-lead ECG from an AD8232 front end (Analog Devices, Wilmington, MA, USA); the AC-coupled impedance signal; the DC-coupled baseline voltage; chest-wall displacement from a chest-strap respiration sensor, used as the respiratory reference; and forehead infrared photoplethysmography (PPG) from a MAX30101 sensor module (Analog Devices, Wilmington, MA, USA) sampled at 400 Hz (Figure 1C).

**Figure 1.**
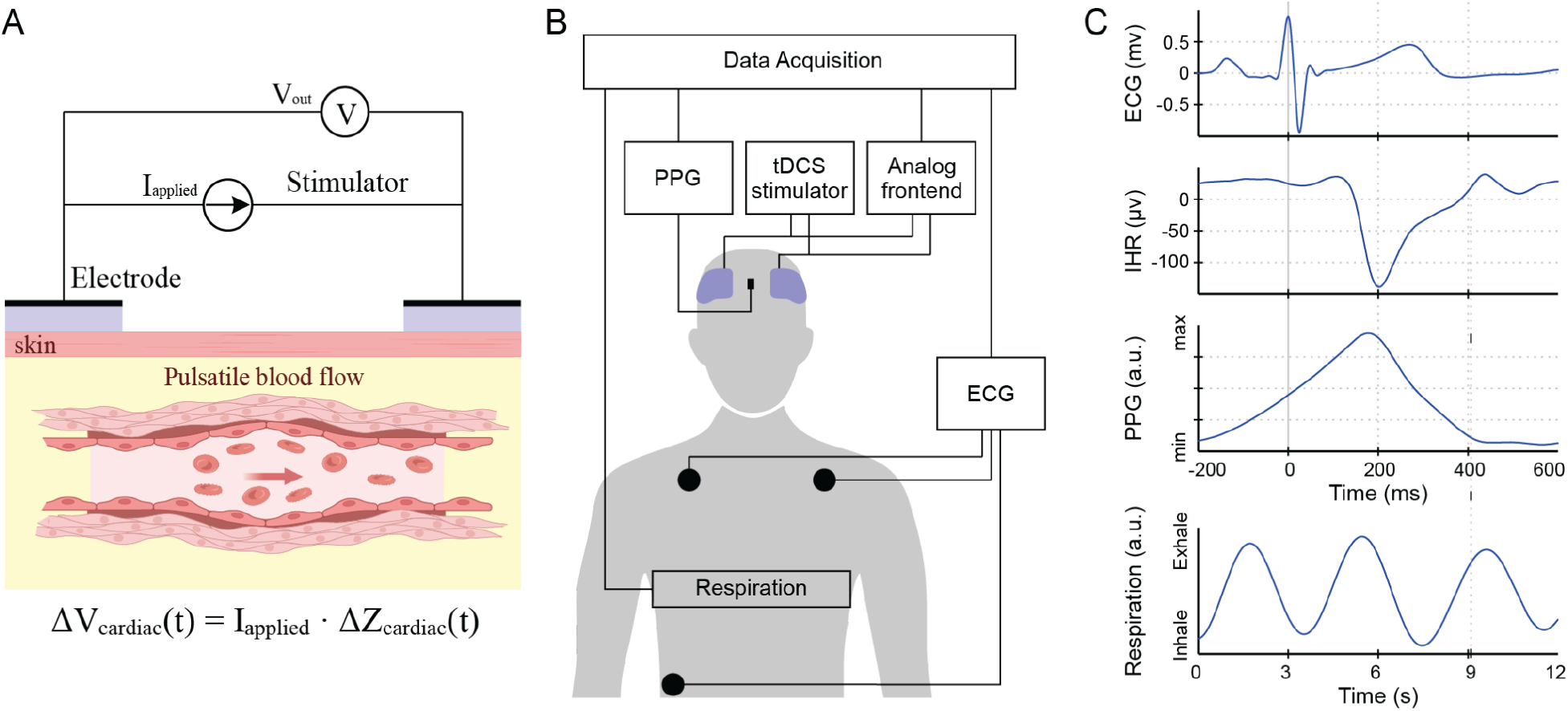
Biophysical basis, experimental setup, and representative cardiac-synchronous signals. (A) System model of constant-current stimulation through time-varying electrode-tissue impedance. Pulsatile blood-volume changes within the stimulation path are hypothesized to contribute a cardiac-impedance term to the total load, which appears at the stimulator output as a cardiac-synchronous voltage fluctuation. (B) Experimental setup. During bifrontotemporal tDCS, a high-impedance analog front-end recorded the stimulator output voltage while ECG, forehead PPG, and respiration signals were acquired for validation. (C) Representative ECG, impedance-derived cardiac (IHR), and forehead PPG waveforms over a single cardiac cycle. Bottom panel: a longer respiration-monitor recording.

For the at-home cohort, stimulation was delivered with a 1 × 1 mini-CT remotely supervised stimulator (Soterix Medical Inc., Woodbridge, NJ, USA), which provides per-session dose control as part of a Remote-Supervised tDCS (RS-tDCS) protocol [18]. The same analog front-end was integrated into a battery-powered standalone logger that recorded the AC-coupled impedance signal at 1 kHz to onboard storage. A Polar H10 chest-strap monitor (Polar Electro, Kempele, Finland), previously validated against ECG for RR-interval and HRV measurement [19], provided the reference RR intervals.

### 2.3 Experimental design

The study included three experiments (Table 1).

**Table 1.** Overview of the Three Experimental Protocols.

| Experiment | Setting / Cohort | Purpose | tDCS protocol | Reference signals | Primary analysis |
| --- | --- | --- | --- | --- | --- |
| Experiment 1 | In-lab/ healthy adults | Feasibility: detect cardiac signal in tDCS output voltage | Continuous 2 mA for ~10 min | ECG, optional respiration/PPG | IHR/IHRV detection and comparison with ECG |
| Experiment 2 | In-lab/ healthy adults | Current dependence | 1-5 mA, 90 s per level, Latin-square order | ECG | Scaling of impedance-derived cardiac signal with current |
| Experiment 3 | At-home/ adults with mild-to-moderate depression | Home validation | 2 mA for 30 min, remotely supervised | Polar H10 RR intervals | HR/HRV validation during home tDCS |

Experiment 1 tested whether a cardiac signal could be recovered from the stimulator output voltage. Participants received 2 mA tDCS for 10 minutes, excluding ramp periods, while ECG and the AC-coupled impedance signal were recorded.

Experiment 2 examined how the impedance-derived signal scaled with applied current. Participants received 1, 2, 3, 4, and 5 mA tDCS in a counterbalanced Latin-square order, with 90 s at each level and ramp transitions between levels. ECG and the AC-coupled impedance signal were recorded throughout.

Experiment 3 evaluated the approach during remotely supervised at-home tDCS. Participants completed a 30-minute session at 2 mA while the AC-coupled impedance signal and Polar H10 RR intervals were recorded.

### 2.4 Signal processing pipeline

All processing used custom Python scripts built on NumPy, SciPy, and NeuroKit2 [20]. The same IHR HR/HRV pipeline was applied to the laboratory and at-home cohorts (Figure 2A), but the cardiac reference differed by setting: single-lead ECG in the laboratory and Polar H10 RR intervals at home. ECG R-peaks were detected with NeuroKit2 [20]. The DC-coupled channel captured the slowly varying baseline output voltage (Figure 2B); subtracting a local baseline estimate (Figure 2C) exposed the low-amplitude pulsatile modulation (Figure 2D).

**Figure 2.**
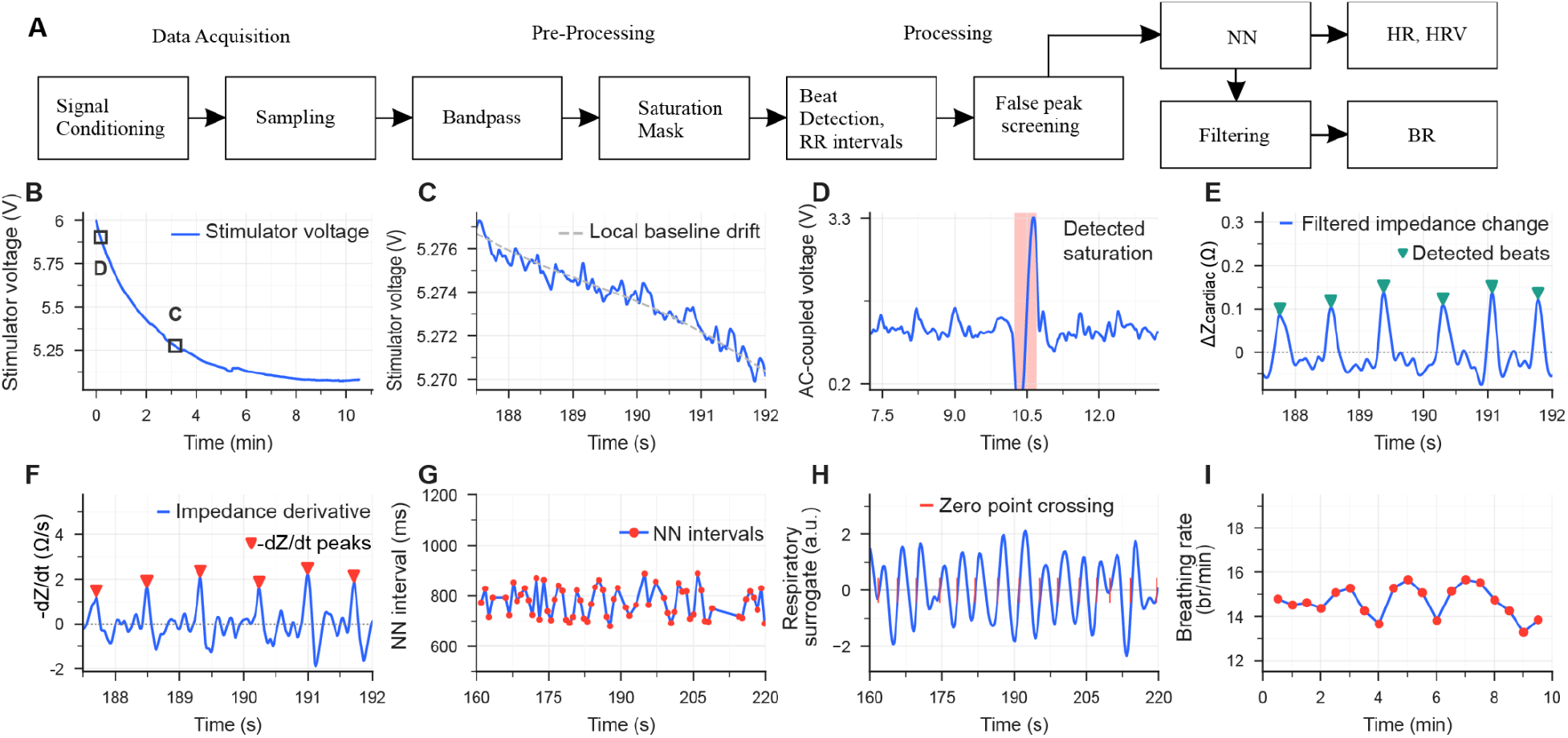
Signal-processing pipeline and representative processing stages for tDCS output impedance-derived measures. (A) Analog signal conditioning and sampling were followed by digital filtering, saturation masking, beat detection, RR-interval screening, and calculation of heart rate, heart rate variability, and respiratory rate. (B) DC-coupled stimulator output during a representative approximately 10-min recording at 2 mA tDCS. Boxes indicate the segments expanded in C and D. (C) Expanded DC-coupled segment showing slow baseline drift with superimposed cardiac fluctuations and the local baseline estimate. (D) Bandpass-filtered AC-coupled segment with the detected saturated region marked for exclusion. (E) Filtered impedance change with detected beats. (F) Impedance derivative with timing-refined fiducials. (G) Accepted normal-to-normal interval tachogram. (H) Impedance-derived respiratory surrogate with detected rising zero crossings. (I) Respiratory-rate trajectory calculated in 60-s windows with a 30-s step.

#### IHR peak detection

Beat-to-beat timing was extracted from the AC-coupled impedance signal with MSPTDfast v2 [21,22]. The sampled 1-kHz waveform was zero-phase bandpass filtered from 0.5 to 5 Hz with a second-order Butterworth filter, and downsampled to 20 Hz for the multi-scale detection stage only; detected beats were then mapped back to the original sampling grid (Figure 2D). Detection used a 30-bpm minimum scale and overlapping 6-s windows with 1-s overlap (Figure 2E) [21]. Each retained beat was relocated to the maximum first derivative of the impedance upstroke (dZ/dt-max), corresponding to the impedance-cardiogram C-point [23], using a Savitzky–Golay derivative with a 41-ms window, evaluated over the preceding 400 ms (Figure 2F).

#### Saturation handling, interval processing, and HRV computation

The AC-coupled impedance signal was screened for amplifier saturation (Figure 2D). Saturation was defined as excursions to ≤0.2 V or ≥3.3 V in laboratory recordings, and to ADC counts ≤10 or ≥4085 in the 12-bit home logger. A symmetric 2.5-s exclusion buffer was applied around each saturated region. Peaks within buffered regions were discarded, and intervals were constructed only between adjacent beats in the same valid region. Intervals outside the 500-1400 ms (43-120 bpm) range were removed. The remaining intervals were screened using a centered 31-interval rolling median, consistent with local-median RR-interval filtering approaches [24,25]; intervals with absolute deviations greater than 15% of the local median were rejected. A subsequent Hampel screen was applied to the absolute successive interval differences using a centered five-difference window and a threshold of the median plus 3.5 scaled median absolute deviations, consistent with established difference-based RR-artifact detection methods [26]. Mean HR, SDNN, and RMSSD [12] were computed from the resulting accepted NN-interval series (Figure 2G). RMSSD differences were retained when the two accepted intervals were originally consecutive and belonged to the same valid region.

#### Respiration processing

In six laboratory participants with a simultaneous chest-respiration reference, respiratory rate was estimated from ECG-derived respiration (EDR) and impedance-derived respiration (IDR). Each NN-interval series was linearly interpolated to 4 Hz, converted to an instantaneous heart-rate tachogram, and transformed into a respiratory surrogate with the van Gent method implemented in NeuroKit2 [8,20]. Rising zero crossings were then detected in the surrogate (Figure 2H). Respiratory rate was calculated in 60-s windows with a 30-s step to produce a breathing-rate trajectory (Figure 2I). The chest reference was bandpass-filtered from 0.1 to 0.4 Hz and evaluated in matching windows.

### 2.5 Validation and statistical analysis

#### In-lab agreement

For the 2 mA block (Experiment 1), HR, SDNN, and RMSSD were each computed once per participant (N = 10). Bias (IHR − ECG), mean absolute error (MAE), and Pearson correlation were calculated across participants.

#### Respiration agreement

Respiratory rate was computed at the participant level in the six-participant subset of the 2 mA block (Experiment 1) that had a respiration-monitor reference. For each respiration estimate (ECG-derived respiration and IDR), the participant-level mean respiratory rate over the session was compared with the respiration-monitor reference (N = 6), and Pearson r, bias, and MAE were reported.

#### At-home agreement

IHR was compared against the Polar H10 reference across 19 at-home sessions from 10 participants. Each metric was computed once per session, and bias, MAE, and Pearson correlation were assessed at the session level for HR, SDNN, and RMSSD.

### 2.6 Frequency-domain and ensemble analyses

Power spectral density (PSD) was estimated with Welch’s method [27] from signals bandpass filtered from 0.5 to 40 Hz. For each participant, the longest continuous portion of the AC-coupled impedance recording during 2 mA tDCS for 10 minutes without detected amplifier saturation was selected automatically, and the ECG was restricted to the same timestamps. PSD estimates used 30-s windows with 50% overlap. The cardiac band was defined as 0.714 to 2.0 Hz, corresponding to 43 to 120 bpm. Dominant ECG and impedance frequencies were compared, and impedance spectral-peak HR was compared with ECG R-peak HR calculated over the same segment. This secondary analysis assessed whether cardiac periodicity could be recovered from the impedance spectrum without beat detection. Spectral-peak analysis provides a dominant average rate but does not resolve the beat-to-beat NN intervals required for HRV computation.

For Experiment 2, ECG-triggered ensemble averages of the AC-coupled impedance signal were calculated separately for each participant and current level from epochs spanning −200 to +800 ms around each ECG R-peak. Before averaging, the signal was bandpass-filtered from 0.5 to 20 Hz with a zero-phase second-order Butterworth filter. This display band was wider than the 0.5–5 Hz band used for beat detection and HRV so that it preserved the sharp trough edges of the impedance pulse while remaining within the analog front-end’s approximately 34 Hz bandwidth.

Three outcomes were derived from each ensemble. Voltage pulse amplitude was defined as the baseline-to-trough excursion: the depth of the dominant trough, identified at 1-ms resolution from 100 to 500 ms after the R-peak, relative to the −200 to −100 ms pre-R baseline.

Within each participant, amplitude was regressed on current across the five current levels. The 10 participant-level slopes were summarized as mean ± standard error and tested against zero with a one-sample t-test. Linearity was summarized by the within-participant coefficient of determination (R^2^) for each five-point fit and reported as the median across participants.

The same within-participant slope analysis and one-sample t-test were applied to current-normalized pulsatile impedance, ΔZ_cardiac_ = ΔV_cardiac_/I_applied_ (mV/mA, equivalent to Ω), and to ECG-to-IHR trough delay, defined as the latency from the ECG R-peak to the dominant trough.

## 3. Results

Our objective was to develop and validate a method for estimating HR, HRV, and respiratory rate using only beat-synchronous oscillations in the tDCS output voltage. We evaluated a hardware and software pipeline in both laboratory subjects and an at-home patient cohort. The AC-coupled output voltage was processed to isolate the pulsatile impedance component, and detected beats were used to construct NN intervals to estimate HR, SDNN, RMSSD, and respiratory rate. We characterized the recovered cardiac-synchronous waveform and compared the impedance-derived estimates with simultaneous ECG, a chest-strap reference, or a respiration monitor, as appropriate.

### 3.1 Recovery of a cardiac-synchronous signal

Removal of the slowly varying tDCS output-voltage baseline, which primarily reflects the constant impedance Z_0_ and slow changes in electrode-skin impedance ΔZ_electrode-skin_(t), revealed a residual pulsatile component (Figure 2B–D). In an exemplary 2 mA recording, this residual exhibits a repeating waveform at the cardiac rate (Figure 3A). Each impedance pulse followed an R-peak in the simultaneous ECG (Figure 3B), and the resulting IHR NN intervals tracked the ECG RR intervals beat by beat (Figure 3C).

**Figure 3.**
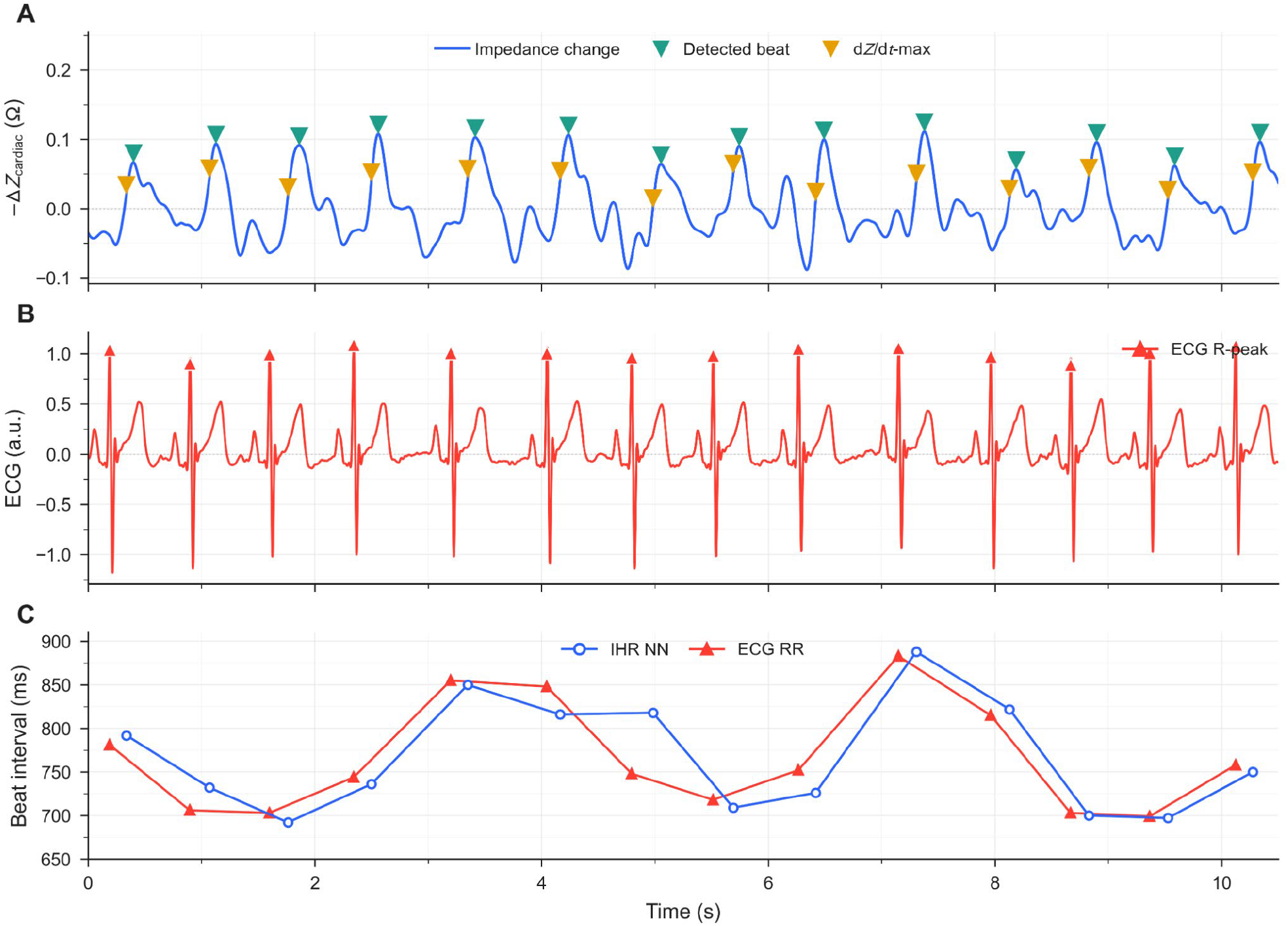
Synchronization of tDCS impedance fluctuations with ECG in a representative 2 mA laboratory recording. (A) Filtered tDCS output impedance over ∼10 s. Repeating impedance pulses (−ΔZ_cardiac_) with reflected cardiac beats (teal triangle peaks) and dZ/dt-max fiducials (amber triangles). (B) Simultaneous ECG with R-peaks (red triangles). (C) Corresponding NN beat intervals based on IHR (blue) and ECG (red).

### 3.2 Current scaling and spectral characteristics

Across 1–5 mA current range, the filtered tDCS output was triggered to the ECG R-peak to characterize the cardiac-synchronous voltage waveform and its dependence on applied current. The averaged voltage shows a cardiac-synchronous signal at every current level in each participant (Figure 4A, representative participant; Supplementary Figure S2, all 10 participants). The waveform included a small deflection approximately 100 ms after the ECG R-peak and a dominant trough at approximately 200 ms. After normalization by applied current, the waveforms represent the associated impedance fluctuation (ΔZ_cardiac_) (Figure 4B).

**Figure 4.**
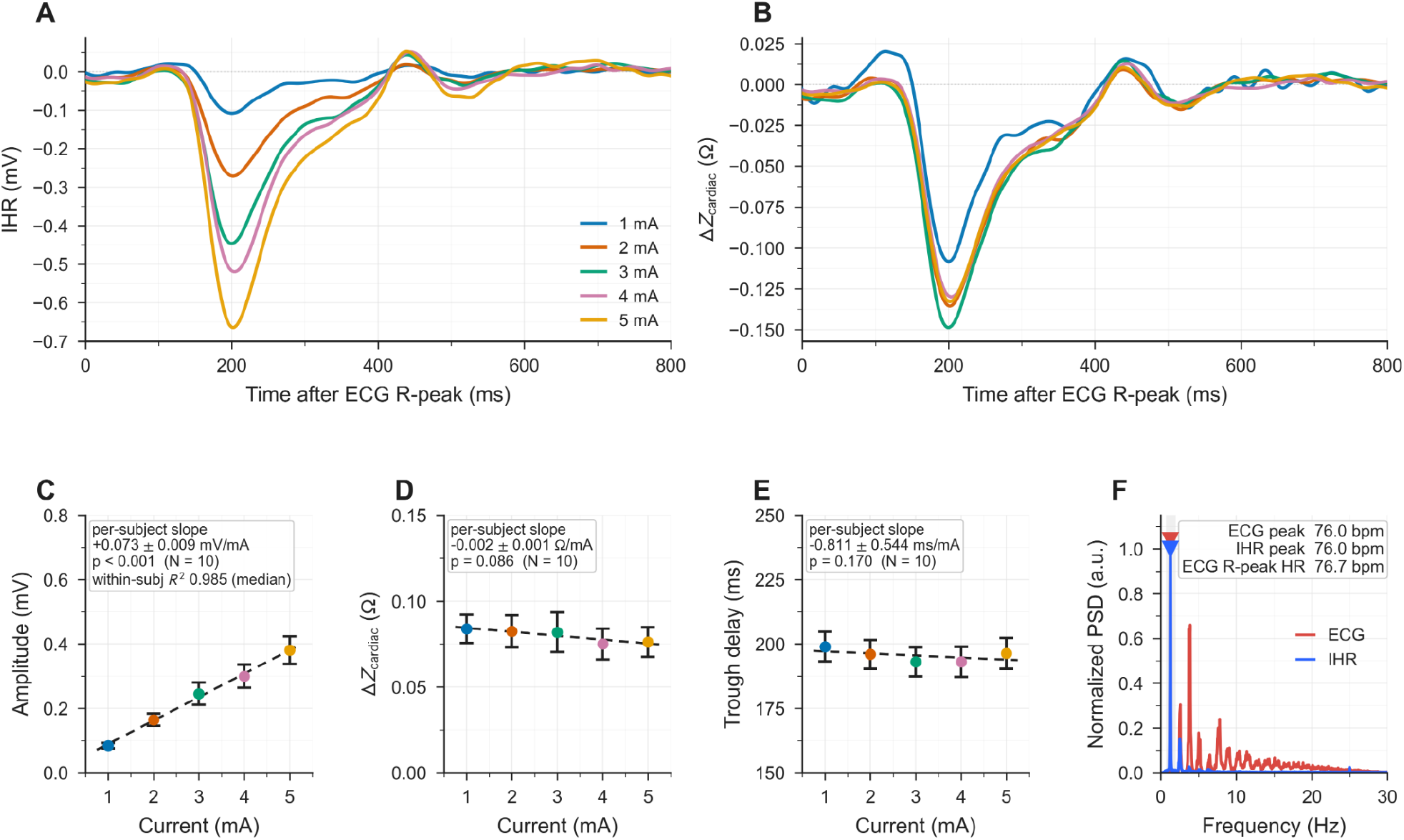
Dependence of IHR amplitude and timing on applied current. (A) Mean ECG R wave-triggered IHR waveform at each current level in a representative participant. (B) Corresponding current-normalized ΔZ_cardiac_ waveforms. (C) Baseline-to-trough voltage amplitude versus current. (D) Current-normalized ΔZ_cardiac_ versus current. (E) ECG-to-IHR trough delay versus current. (F) Representative Welch PSDs of ECG and IHR; triangles mark the dominant frequencies, and the inset reports spectral-peak rates and time-domain ECG R-peak HR. Points in C–E are means across participants at each current level (± SEM across participants); dashed lines are least-squares trends through those points, which for this balanced design (every participant measured at all five levels) equal the mean per-participant slope. The boxed slope ± SE and p value are computed from the 10 per-participant slopes (N = 10 participants); C also reports the median within-participant R^2^.

Cardiac voltage amplitude, measured from baseline to trough, increased linearly with applied current. The mean slope across the 10 participant-specific regressions was 0.073 ± 0.009 mV/mA (mean ± SE; p < 0.001; Figure 4C). Within-participant coefficients of determination ranged from 0.859 to 0.999, with a median R^2^ of 0.985. After division by applied current, ΔZ_cardiac_ was not significantly associated with current intensity (mean participant-level slope ± SE, −0.002 ± 0.001 Ω/mA; p = 0.086; Figure 4D). ΔZ_cardiac_ was also not significantly associated with the mean DC output voltage during each 90-s current block (Supplementary Figure S3).

Pooling all 50 participant–current measurements obtained from 10 participants at five current levels, ΔZ_cardiac_ averaged 0.080 ± 0.029 Ω (mean ± SD; Figure 4D), with individual measurements ranging from 0.020 to 0.149 Ω. Averaging across participants, mean ΔZ_cardiac_ at each current level ranged from 0.075 to 0.084 Ω (SD across the five current-level means, 0.004 Ω). Averaging across current levels, mean ΔZcardiac for each participant ranged from 0.029 to 0.131 Ω (SD across the 10 participant means, 0.029 Ω). Thus, variability was greater between participants than across current levels (Supplementary Figure S3).

The delay from the ECG R-peak to the IHR trough was not significantly associated with current intensity (mean participant-level slope ± SE, −0.81 ± 0.54 ms/mA; p = 0.170; N = 10; Figure 4E). Across the 50 participant-current observations, the delay averaged 196 ± 18 ms (mean ± SD).

The ECG and the cardiac-synchronous impedance signal had dominant spectral peaks within the cardiac band (Figure 4F). Across participants, the peaks occurred in the same Welch frequency bin in 8 of 10 participants and within one bin in all 10 participants (bin width, 0.033 Hz, equivalent to 2 bpm; Supplementary Figure S4). Over the continuous segments without detected amplifier saturation defined in Section 2.6, impedance spectral-peak HR differed from ECG R-peak HR by a bias of −0.18 bpm and an MAE of 0.88 bpm (r = 0.992; N = 10). Individual errors ranged from −1.56 to +2.22 bpm, with 8 of 10 participants within 1.50 bpm. This secondary analysis confirmed cardiac periodicity in the impedance signal. Since this analysis cannot resolve beat-to-beat timing variations, all primary HR and HRV estimates were obtained using the time-domain beat-based pipeline described in Section 2.4.

### 3.3 In-lab validation against ECG and a respiration reference

In the laboratory cohort, cardiac and respiratory measures derived from impedance fluctuations during 2 mA tDCS were validated against ECG and respiration references. Impedance-derived fiducials produced NN intervals that tracked the simultaneous ECG RR intervals beat by beat (representative subject, Figure 5A and B). In the same subject, IDR (from the impedance NN intervals) and EDR (from the ECG RR intervals) both followed the respiration-monitor reference as respiratory waveforms (Figure 5C) and breathing-rate trajectories (Figure 5D).

**Figure 5.**
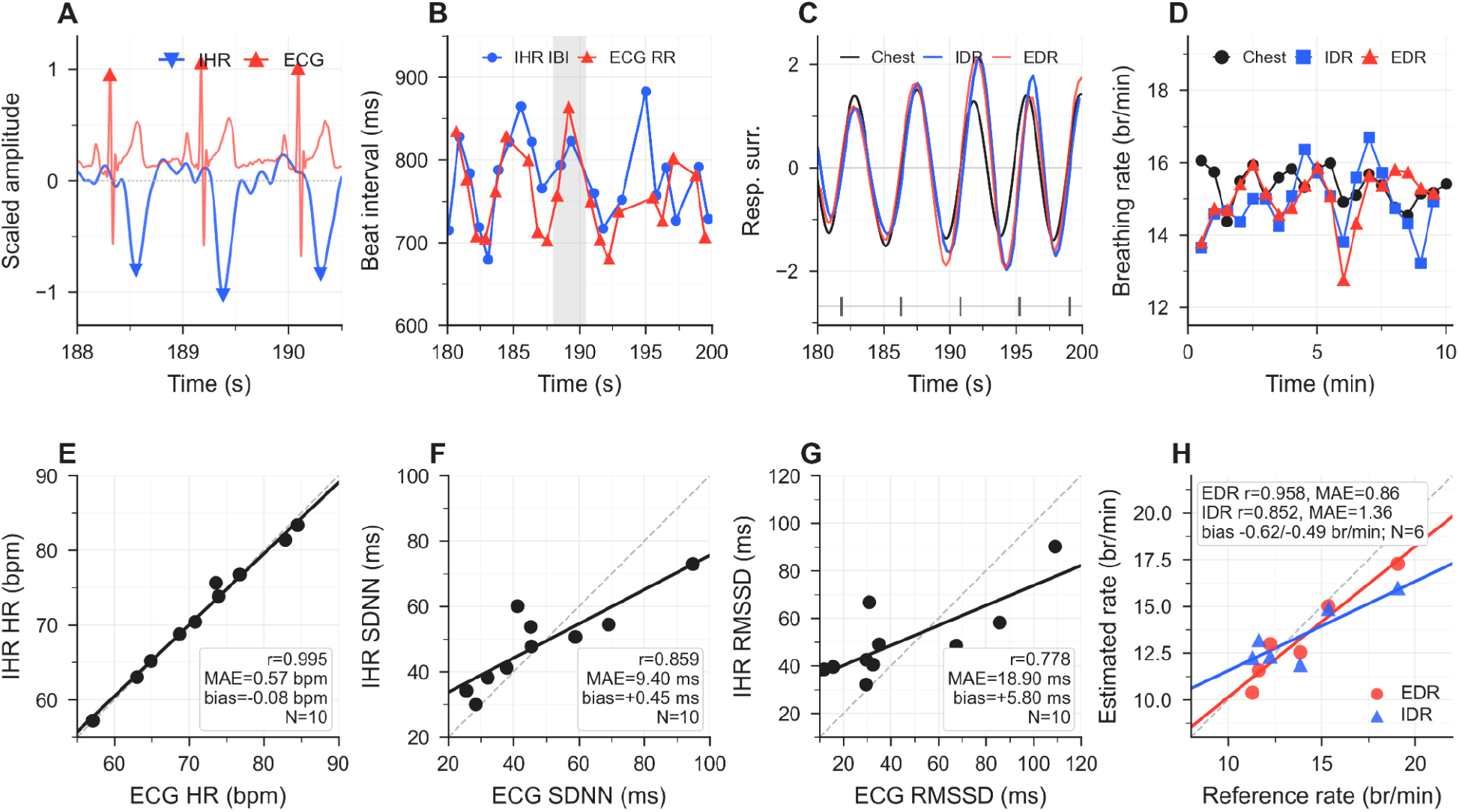
In-lab validation of impedance-derived HR, HRV, and respiration during 2 mA tDCS. Panels A– D are from a representative laboratory recording, displayed over different time windows. (A) Representative IHR and ECG waveforms with fiducials. (B) Corresponding intervals; shading marks the window expanded in A. (C) Respiration-monitor reference, IDR, and ECG-derived respiration surrogates; ticks mark the IDR rising zero crossings counted for rate. (D) Corresponding breathing-rate trajectories over the ∼10-min block. (E–G) Participant-level HR, SDNN, and RMSSD compared with ECG (N = 10). (H) Participant-mean respiratory rate versus the respiration-monitor reference (N = 6); bias is reported for EDR then IDR. In E–H, each point is one participant: the dashed diagonal marks perfect agreement and the solid line is the least-squares fit to the points. Annotations give Pearson r, MAE, and bias (estimate − reference). EDR, ECG-derived respiration; IDR, impedance-derived respiration.

Across the 10 participants, mean IHR bias was −0.08 bpm with an MAE of 0.57 bpm relative to ECG. Participant-level IHR and ECG HR values were correlated at r = 0.995 (Figure 5E). Individual HR errors ranged from −1.50 to +2.05 bpm. Nine participants had absolute errors of 1.50 bpm or less, and all 10 had absolute errors of 2.05 bpm or less (Table 2).

**Table 2.** Participant-level results for ECG-referenced IHR validation.

| ID | ECG HR (bpm) | IHR HR (bpm) | HR error (bpm) | ECG SDNN (ms) | IHR SDNN (ms) | SDNN error (ms) | ECG RMSSD (ms) | IHR RMSSD (ms) | RMSSD error (ms) | Clean RR (n) |
| --- | --- | --- | --- | --- | --- | --- | --- | --- | --- | --- |
| S1 | 64.90 | 65.19 | +0.29 | 32.10 | 38.16 | +6.06 | 15.72 | 39.48 | +23.76 | 507 |
| S2 | 84.53 | 83.36 | -1.17 | 45.43 | 47.71 | +2.28 | 32.47 | 40.47 | +8.00 | 224 |
| S3 | 70.82 | 70.42 | -0.40 | 37.91 | 41.19 | +3.28 | 29.63 | 42.48 | +12.85 | 436 |
| S4 | 76.80 | 76.75 | -0.05 | 25.52 | 34.19 | +8.67 | 11.82 | 38.47 | +26.65 | 931 |
| S5 | 63.04 | 63.02 | -0.02 | 45.23 | 53.72 | +8.49 | 35.03 | 48.94 | +13.91 | 425 |
| S6 | 82.91 | 81.41 | -1.50 | 69.03 | 54.42 | -14.61 | 85.72 | 58.17 | -27.55 | 93 |
| S7 | 57.12 | 57.20 | +0.08 | 41.18 | 60.04 | +18.86 | 30.89 | 66.70 | +35.81 | 420 |
| S8 | 73.59 | 75.64 | +2.05 | 94.80 | 72.94 | -21.86 | 109.08 | 90.16 | -18.92 | 472 |
| S9 | 73.94 | 73.84 | -0.10 | 28.41 | 29.99 | +1.58 | 29.57 | 32.09 | +2.52 | 652 |
| S10 | 68.73 | 68.78 | +0.05 | 58.89 | 50.62 | -8.27 | 67.51 | 48.45 | -19.06 | 546 |

Note. Error columns are IHR – ECG. Clean IHR RR is the final accepted interval count.

IHR-derived SDNN had a mean bias of +0.45 ms and an MAE of 9.40 ms relative to ECG, with *r* = 0.859 (Figure 5F). IHR-derived RMSSD had a mean bias of +5.80 ms and an MAE of 18.90 ms, with *r* = 0.778 (Figure 5G). Participant-level HR, SDNN, and RMSSD values are provided in Table 2, with cohort-level statistics in Table 3.

**Table 3.** Cohort-level summary statistics for ECG-referenced IHR validation (Experiment 1, 2 mA block, N = 10). Reports mean ± SD, bias, MAE, and Pearson r for HR (bpm), SDNN (ms), and RMSSD (ms).

| Metric | ECG reference<br>mean ± SD | IHR mean ±<br>SD | Bias (IHR –<br>ECG) | MAE | Pearson r | N |
| --- | --- | --- | --- | --- | --- | --- |
| HR (bpm) | 71.64 ± 8.62 | 71.56 ± 8.26 | -0.08 | 0.57 | 0.995 | 10 |
| SDNN (ms) | 47.85 ± 21.23 | 48.30 ± 12.93 | +0.45 | 9.40 | 0.859 | 10 |
| RMSSD (ms) | 44.74 ± 31.90 | 50.54 ± 17.23 | +5.80 | 18.90 | 0.778 | 10 |

Note. Bias is IHR – ECG. MAE is the mean absolute error across participants.

Across the six participants with a simultaneous respiration-monitor reference, participant-mean respiratory rate from IDR showed a bias of −0.49 breaths/min, MAE of 1.36 breaths/min, and r = 0.852. ECG-derived respiration bias was −0.62 breaths/min, with an MAE of 0.86 breaths/min, and r = 0.958 (Figure 5H).

### 3.4 At-home validation against Polar H10

Across 19 at-home 30-minute stimulation sessions from 10 participants, IHR-derived HR had a mean bias of −1.43 bpm and an MAE of 1.43 bpm relative to HR calculated from Polar H10 RR intervals. Session-level IHR and Polar H10 HR values were correlated at r = 0.995 (Figure 6A).

**Figure 6.**
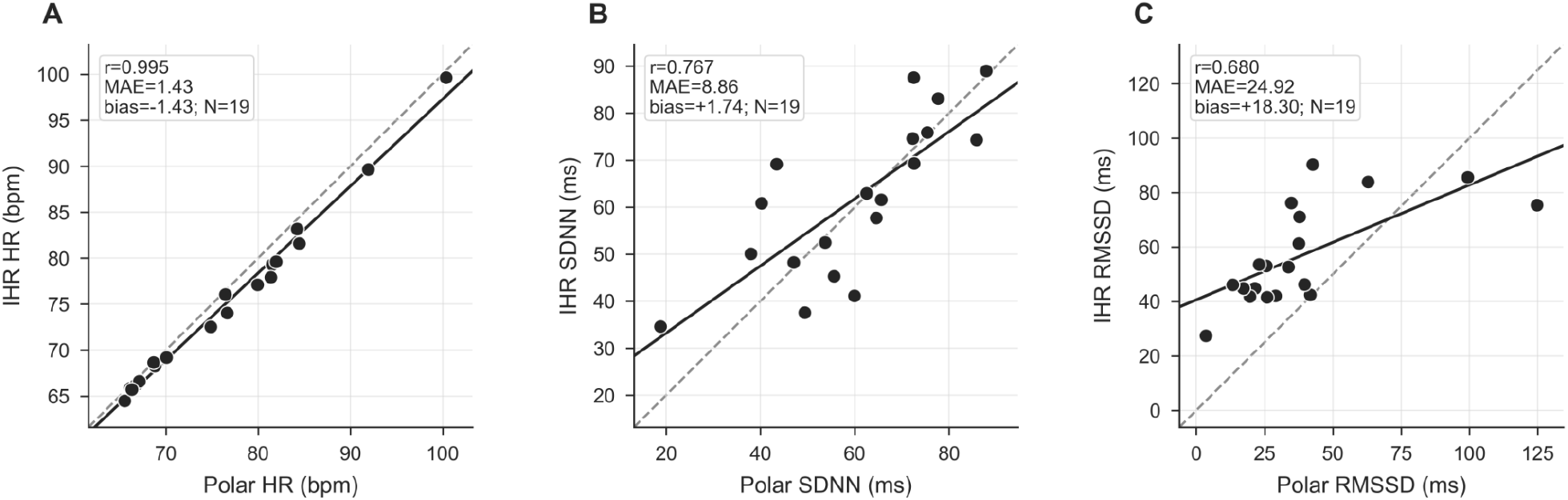
At-home validation against the Polar H10 reference (Experiment 3; 19 sessions from 10 participants). (A) HR, (B) SDNN, and (C) RMSSD. Each point is one session, so participants contributing multiple sessions appear more than once; the dashed diagonal marks perfect agreement and the solid line is the least-squares fit to the points. Annotations give Pearson r, MAE, and bias (IHR − Polar H10).

IHR-derived SDNN had a mean bias of +1.74 ms and an MAE of 8.86 ms relative to the Polar H10 reference, with *r* = 0.767 (Figure 6B). IHR-derived RMSSD mean bias was +18.30 ms with an MAE of 24.92 ms, and *r* = 0.680 (Figure 6C).

Session-level results are provided in Table 4. Bland-Altman analyses at the session and participant levels are provided in Supplementary Figures S5 and S6, respectively. When repeated session metrics were averaged within each participant, the MAEs were 1.50 bpm for HR, 7.73 ms for SDNN, and 27.42 ms for RMSSD (Supplementary Figure S6).

**Table 4.** At-home validation against the Polar H10 reference (Experiment 3; 19 sessions across 10 participants).

| Metric | Polar<br>H10 ref<br>mean ±<br>SD | IHR<br>mean ±<br>SD | Bias (IHR<br>– Polar<br>H10) | MAE | Pearson r |
| --- | --- | --- | --- | --- | --- |
| HR (bpm) | 76.47 ±<br>9.72 | 75.04 ±<br>9.24 | -1.43 | 1.43 | 0.995 |
| SDNN<br>(ms) | 60.11 ±<br>17.79 | 61.86 ±<br>16.56 | +1.74 | 8.86 | 0.767 |
| RMSSD<br>(ms) | 38.56 ±<br>29.25 | 56.85 ±<br>18.15 | +18.30 | 24.92 | 0.680 |

Note. Bias is IHR – Polar H10. MAE is the mean absolute error. Values are session-level summaries for the at-home cohort.

## 4. Discussion

We demonstrate that cardiac and respiratory metrics can be derived from voltage fluctuations recorded at tDCS device output without additional electrodes or sensors. In the laboratory cohort, impedance-derived heart rate differed from simultaneous ECG by a mean absolute error of 0.57 bpm, while the mean absolute errors for SDNN and RMSSD were 9.40 ms and 18.90 ms, respectively. Impedance-derived respiratory rate differed from the respiration-monitor reference by 1.36 breaths/min. Across 19 at-home stimulation sessions from 10 participants, the mean absolute errors relative to values calculated from Polar H10 RR intervals were 1.43 bpm for heart rate, 8.86 ms for SDNN, and 24.92 ms for RMSSD. Together, these findings support physiological monitoring (heart rate, heart rate variability, and respiration) using only the tDCS output voltage.

### Biophysical basis

The recovered cardiac-synchronous signal is consistent with the principles of impedance plethysmography and impedance cardiography [4,5]. During constant-current tDCS, heartbeat-related changes in stimulation-path impedance produce corresponding output-voltage fluctuations that increase with applied current. This scaling is consistent with an impedance-derived origin of the recovered waveform. Across 50 participant-current measurements from 10 participants tested at five current levels, mean current-normalized pulsatile impedance (ΔZ_cardiac_=ΔV_cardiac_/I_applied_) was 0.080 ± 0.029 Ω (mean ± SD). The pulsatile impedance change ΔZ_cardiac_ was not significantly associated with current intensity (mean participant-level slope, −0.002 Ω/mA; p = 0.086). The latency from the ECG R-peak to the ΔZ_cardiac_ trough was also not significantly associated with current intensity and is consistent with cardiac pre-ejection and pulse-transit time [28].

### Accuracy Considerations

Mean HR depends on the average beat-to-beat interval across a recording, so small beat-timing errors largely average out. In contrast, SDNN and especially RMSSD depend on variability among individual intervals, making them more sensitive to timing uncertainty. The observed biases followed this expected pattern. SDNN was nearly unbiased in both cohorts (+0.45 ms in the laboratory and +1.74 ms at home), whereas RMSSD was systematically overestimated (+5.80 and +18.30 ms, respectively), consistent with fiducial-placement jitter, which disproportionately inflates successive interval differences. Contributing factors include beat-to-beat variation in pulse transit time [28,29] and uncertainty in identifying a consistent fiducial point on the broader impedance upstroke [23]. Similar limitations affect pulse-based methods, including photoplethysmography (PPG), in which physiological pulse-transit variability and fiducial uncertainty constrain HRV accuracy relative to ECG.

### Applications and generalization

The approach developed here (Figure 2A) may extend to other invasive and noninvasive electrical stimulation techniques, enabling onboard physiological monitoring without additional sensors. Implementation would require waveform-specific acquisition and signal processing, including demodulation for AC and pulsed stimulation.

Beyond cardiac and respiratory signals, the stimulator output voltage may contain additional endogenous and impedance-related information, including ocular activity, myogenic activity, cortical electrical activity, and slower impedance changes associated with blood volume, respiration, motion, and the electrode-skin interface. These signals were not studied here but can be following analogous principles.

## CRediT authorship contribution statement

**Yahia Abdalla:** Methodology, Software, Formal analysis, Investigation, Visualization, Writing – original draft, Writing – review & editing. **Benjamin Babaev:** Investigation, Formal analysis, Visualization. **Kevin Walsh:** Investigation. **Catarina Ferraz:** Investigation. **Leigh Charvet:** Conceptualization, Resources, Funding acquisition, Supervision. **Giuseppina Pilloni:** Conceptualization, Resources, Funding acquisition, Supervision. **Marom Bikson:** Conceptualization, Supervision, Funding acquisition, Resources, Writing – original draft, Writing – review & editing. **Mohamad FallahRad:** Conceptualization, Supervision, Funding acquisition, Resources, Writing – original draft, Writing – review & editing.

