## Supplementary Figures S1-S6 for "Impedance-Derived Heart Rate and Heart Rate Variability from tDCS Output Voltage: Sensorless Physiological Monitoring During Electrical Neuromodulation"

### Supplementary Material

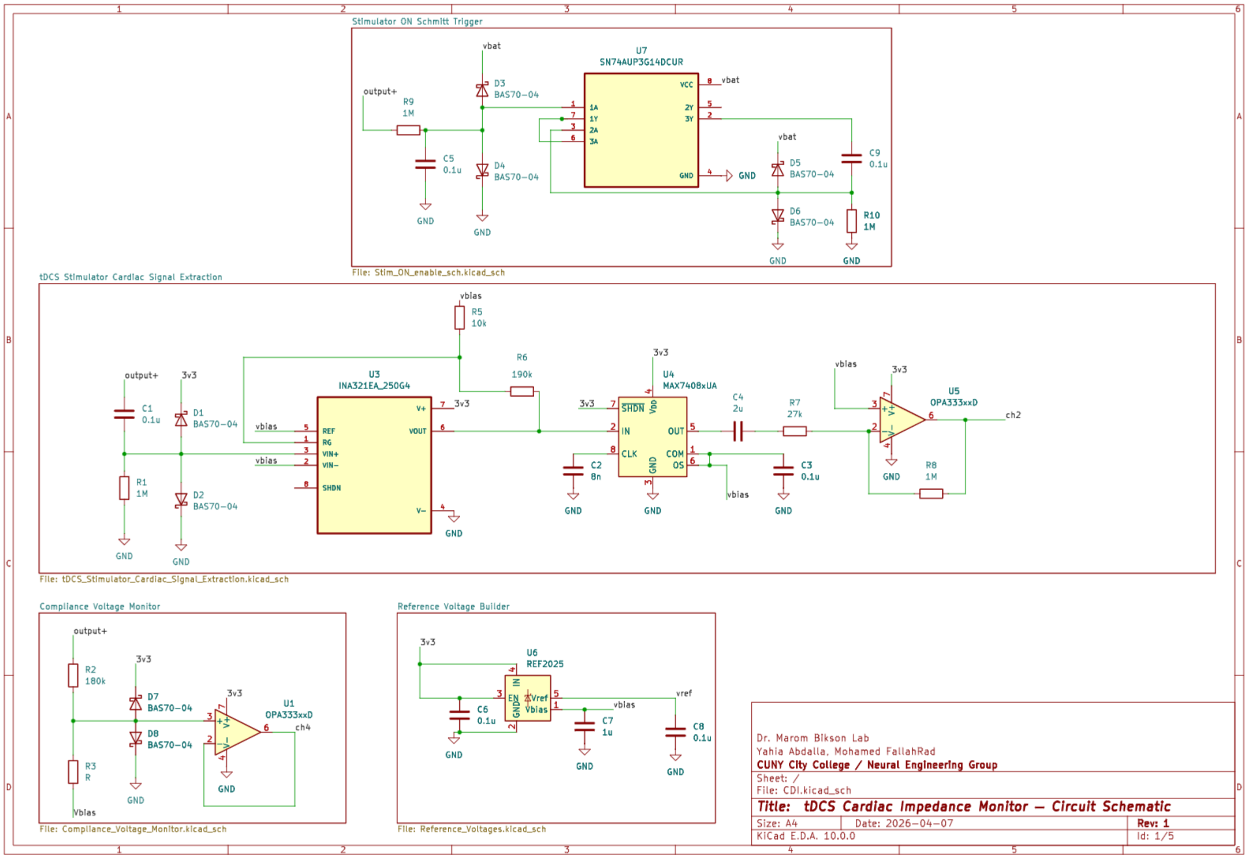


### Supplementary Figure S1. Analog front-end circuit schematic.

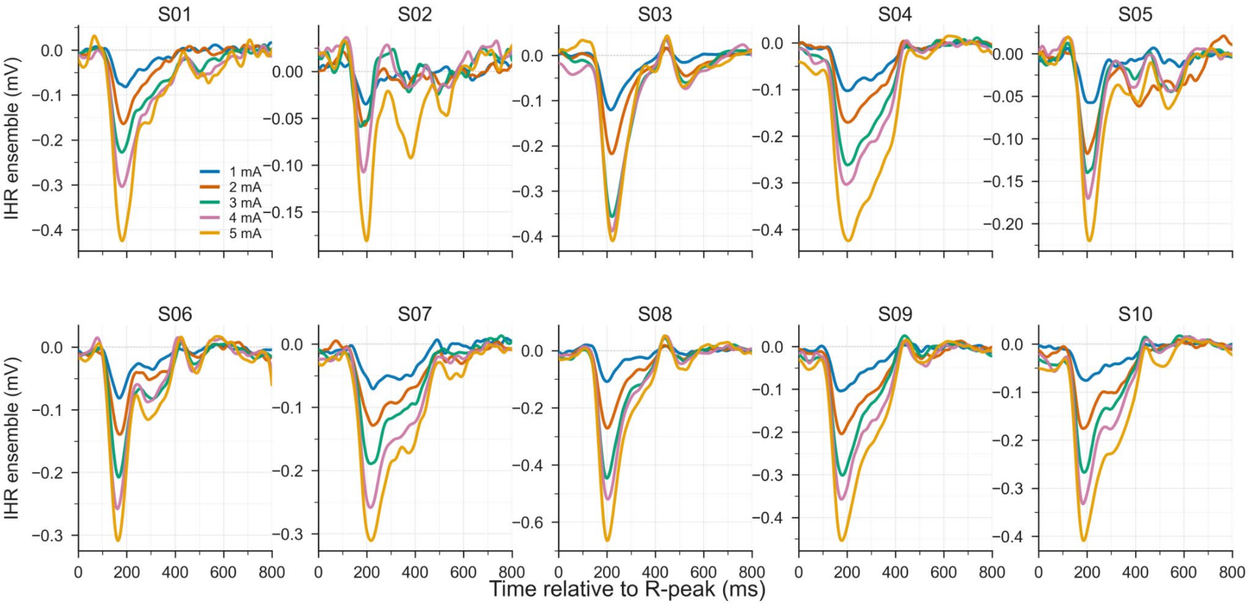

**Supplementary Figure S2.** Participant-level ECG-triggered ensemble averages across the 1–5 mA current sweep. The figure includes all 10 in-lab participants in Experiment 2 (S01–S10). Each panel shows the mean ensemble waveform at each current level on a common time axis (−200 to +800 ms relative to the ECG R-peak).

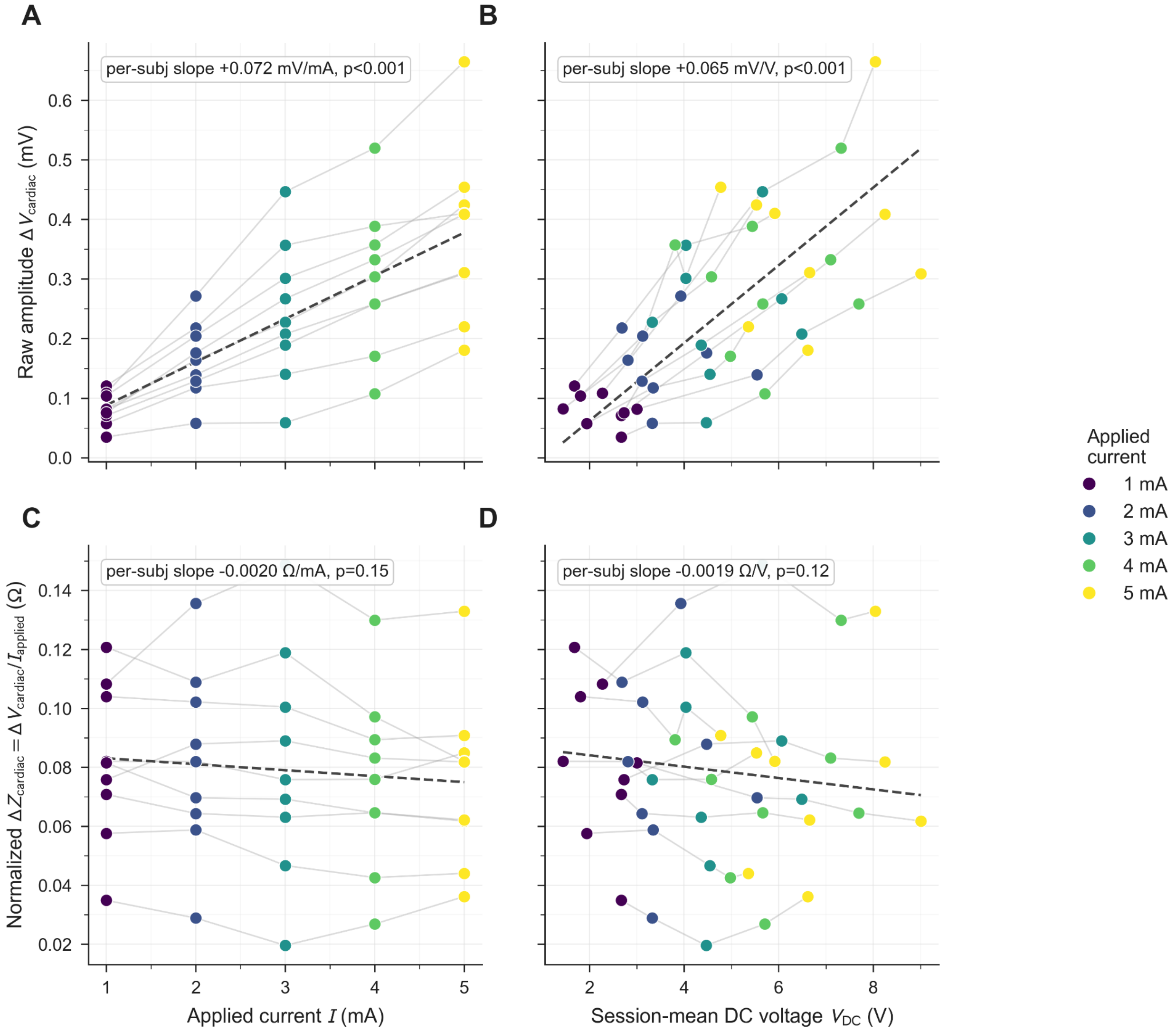


**Supplementary Figure S3.**
Determinants of IHR pulse amplitude. Nine of the ten Experiment 2 participants had valid DC voltage recordings, yielding 45 participant-current observations (9 participants × 5 current levels); points are colored by applied current. Because all panels are restricted to this nine-participant subset, the slopes and p values differ slightly from the ten-participant analysis reported in Section 3.2. (A, B) Raw baseline-to-trough amplitude ($\Delta V_{cardiac}$) increased with both applied current (A; mean participant-level slope, +0.072 mV/mA; $p < 0.001$) and session-mean DC output voltage (B; +0.065 mV/V; $p < 0.001$). These predictors are collinear because VDC = $I_{applied} \cdot Z_0$. (C, D) After normalization by applied current, $\Delta Z_{cardiac}(\Omega)$ showed no significant association with current (C; mean participant-level slope, −0.002 Ω/mA; $p = 0.15$) or DC voltage (D; −0.002 Ω/V; $p = 0.12$). This normalization removes the current scaling predicted by Eq. (4). Individual $\Delta Z_{cardiac}$ observations span 0.020–0.149 Ω, a spread that is almost entirely between participants

(participant means, 0.029–0.131 Ω) rather than across current levels. Dashed lines show the mean per-participant slope.

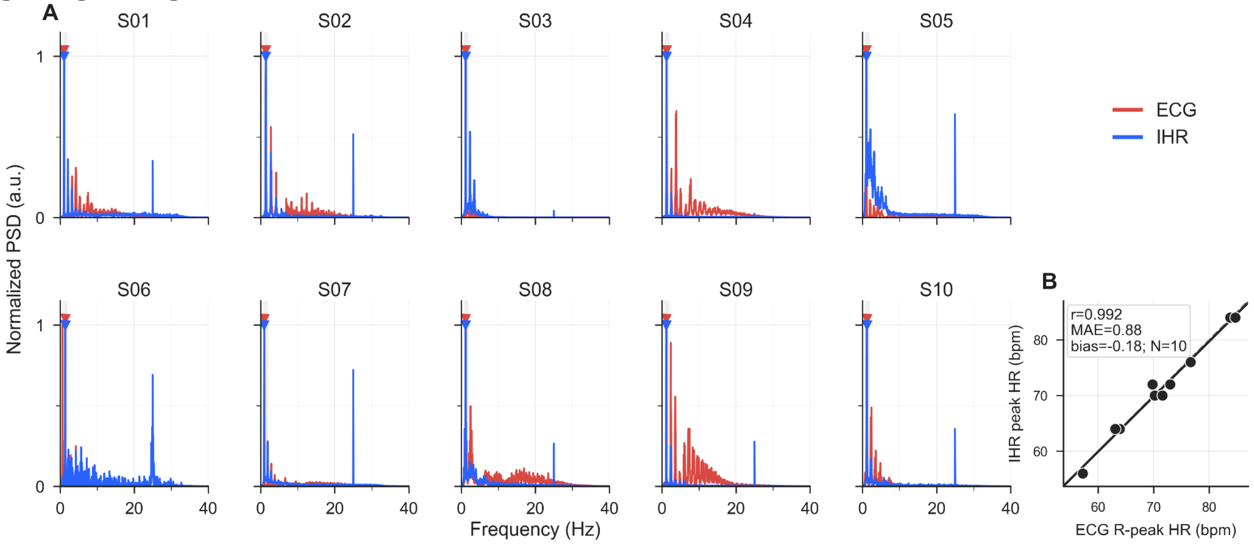


**Supplementary Figure S4.** Frequency-domain cardiac summary for Experiment 1 (N = 10). (A) Per-participant Welch PSD overlays of ECG and the AC-coupled impedance signal over each participant's longest continuous segment without detected amplifier saturation; ECG was restricted to the same timestamps. ECG and impedance peaks occurred in the same frequency bin in 8 of 10 participants and within one bin in all 10 participants. (B) Impedance spectral-peak HR versus ECG R-peak HR over the same segment; each point is one participant. The dashed diagonal marks perfect agreement and the solid line is the least-squares fit (MAE = 0.88 bpm, r = 0.992). Individual errors ranged from −1.56 to +2.22 bpm, with 8 of 10 participants within 1.50 bpm. Because the reference is recomputed over the matched segment, it differs slightly from the full-recording ECG HR in Table 2. Dashed line, identity; solid line, least-squares fit.

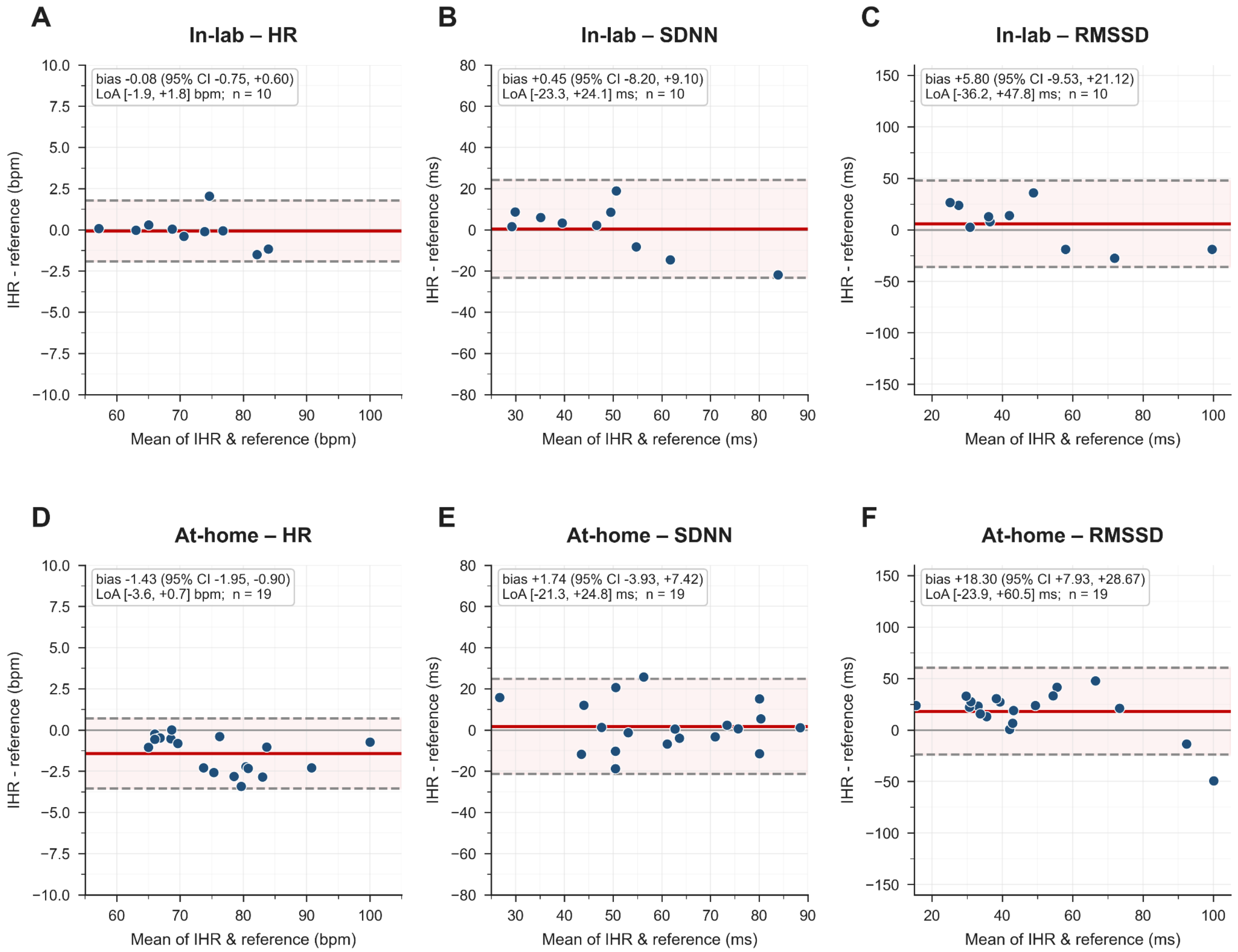


**Supplementary Figure S5.** Bland–Altman agreement for impedance-derived HR, SDNN, and RMSSD. Top row (A–C): in-lab cohort versus simultaneous ECG (N = 10). Bottom row (D–F): at-home cohort versus Polar H10 (19 sessions from 10 participants). Each point is one participant in A–C and one session in D–F. Each panel plots the difference (IHR − reference) against the mean of the two methods; the solid line is the mean difference (bias), the dashed lines are the 95% limits of agreement (bias ± 1.96 SD of the differences), and the bias with its 95% confidence interval is annotated within each panel.

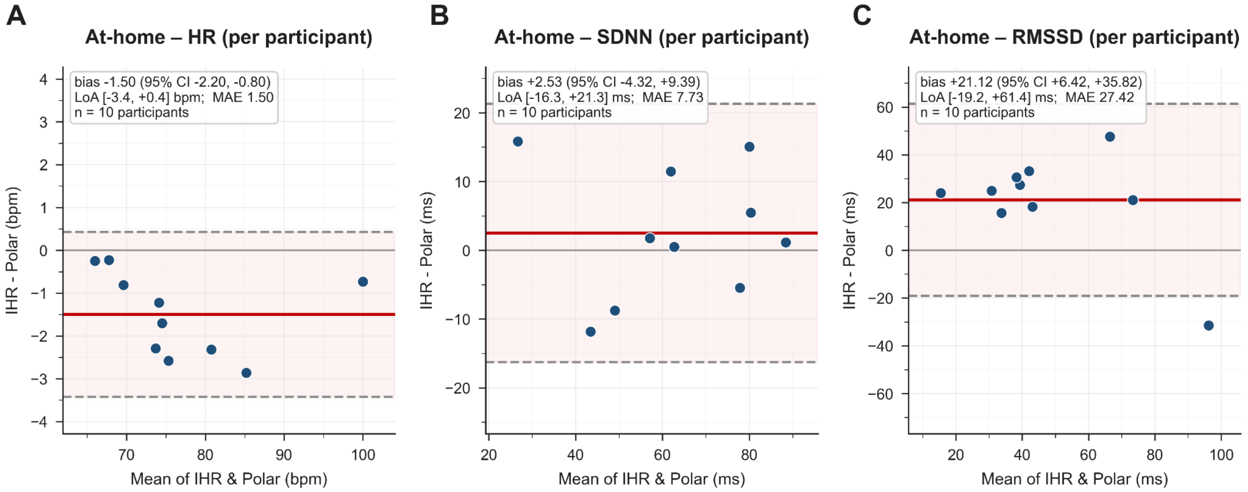


**Supplementary Figure S6**. Participant-level Bland–Altman agreement for the at-home cohort. The 19 at-home sessions come from 10 participants; each participant is collapsed to a single point (the mean of that participant's session-level IHR and Polar H10 values) before Bland–Altman agreement is computed on the 10 participant means. (A) HR, (B) SDNN, (C) RMSSD. Each point is one participant; the solid line is the mean difference (bias) and the dashed lines are the 95% limits of agreement (bias ± 1.96 SD of the differences), with the bias and its 95% confidence interval annotated in each panel. Participant-level mean absolute errors were 1.50 bpm (HR), 7.73 ms (SDNN), and 27.42 ms (RMSSD), compared with 1.43 bpm, 8.86 ms, and 24.92 ms at the session level. This participant-level view avoids treating repeated sessions as independent participant-level observations in the session-level analysis (Supplementary Figure S5) at the cost of power (n = 10).
